# Rapid resistome mapping using nanopore sequencing

**DOI:** 10.1101/067652

**Authors:** Eric van der Helm, Lejla Imamovic, Mostafa M Hashim Ellabaan, Willem van Schaik, Anna Koza, Morten O. A. Sommer

**Affiliations:** Novo Nordisk Foundation Center for Biosustainability, Technical University of Denmark, DK-2870 Hørsholm, Denmark; Department of Medical Microbiology, University Medical Center Utrecht, Utrecht, The Netherlands

**Keywords:** Nanopore sequencing, MinION, resistome, functional metagenomics

## Abstract

The emergence of antibiotic resistance in human pathogens has become a major threat to modern medicine and in particular hospitalized patients. The outcome of antibiotic treatment can be affected by the composition of the gut resistome either by enabling resistance gene acquisition of infecting pathogens or by modulating the collateral effects of antibiotic treatment on the commensal microbiome. Accordingly, knowledge of the gut resistome composition could enable more effective and individualized treatment of bacterial infections. Yet, rapid workflows for resistome characterization are lacking. To address this challenge we developed the poreFUME workflow that deploys functional metagenomic selections and nanopore sequencing to resistome mapping. We demonstrate the approach by functionally characterizing the gut resistome of an ICU patient. The accuracy of the poreFUME pipeline is >97 % sufficient for the reliable annotation of antibiotic resistance genes. The poreFUME pipeline provides a promising approach for efficient resistome profiling that could inform antibiotic treatment decisions in the future.

## Background

It is estimated that every year 700,000 people die of resistant infections [1]. Antibiotic resistance by human pathogens has become a major threat, in particular for hospitalized patients [1, 2]. Bacterial infections by resistant pathogens are also coupled with an increase in healthcare costs[3]. The gut microbiome comprises a diverse set of antibiotic resistance genes which may impact antibiotic treatment outcomes in at least two ways[4-6]. First, the gut microbiome can act as a reservoir of resistance genes that can be acquired by infecting human pathogens leading to evolution of resistance during infection. Indeed, a close evolutionary relationship between resistance genes in pathogens and commensals has been found [7]. Second, the gut resistome impacts the extent to which the commensal microbiota is affected by antibiotic treatment. Studies of preterm infants and their response to antibiotic treatment suggest that the collateral damage to the commensal microbiota can be predicted from the resistome status at the start of treatment[8]. Accordingly, there is an increasing interest in the development of clinically applicable workflows that enable expedited and comprehensive characterization of the gut resistome. Unfortunately, given the diversity of antibiotic resistance genes in the gut microbiota, sequencing based methods alone cannot enable a representative characterization of the gut resistome. Instead, functional metagenomic selections, which circumvent the culturing step of individual gut microbes, allow less biased interrogation of the gut resistome[9]. Consequently, rapid resistome profiling using a functional-metagenomic approach would be a viable approach to guide personalized antibiotic treatment.

A functional metagenomic workflow consists of several steps, of which the final step is the analysis of metagenomic sequencing data (Figure 1)[10]. Traditionally, sequencing data was obtained using Sanger sequencing, [11, 12], yet, other high-throughput sequencing technologies such as 454 pyrosequencing [13] and Illumina sequencing [6, 14] have been applied to analyze functional metagenomic assays as well. The PARFuMS pipeline based on Illumina data was used to profile the antibiotic resistome of soil and of the human microbiome[4, 6, 14], and PacBio SMRT data has been used to sequence large metagenomic insert libraries (~40 kb) from fosmids [15]. There are several challenges related to data processing and annotation in functional metagenomic selections. For Sanger sequencing, the data annotation is usually done on non-complete contigs. In each Sanger set, contigs can be closed by using primer walking. However, primer walking is hardly feasible for high-throughputs datasets and requires weeks to complete. Short read sequencing based on the Illumina platform offers a high-throughput method, yet, contig assembly can be hampered by repetitive sequences in the original insert. A workflow based on PacBio SMRT data circumvents such assembly challenges; however, this technology has a significant capital cost requirement, a large laboratory footprint and is technically demanding limiting point of care applications [16]. In contrast, nanopore sequencing may be able to address these challenges enabling on site monitoring of resistomes in both clinical and environmental settings.

**Figure 1:**
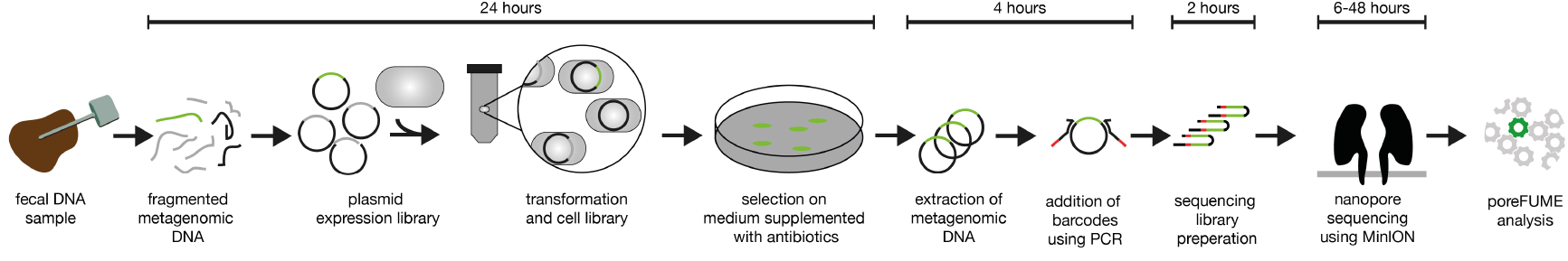
Overview of the poreFUME workflow consisting of the construction and sequencing of a metagenomic library. Fecal DNA is extracted, purified, fragmented and cloned into a shuttle vector. The library of plasmids is transformed in an *E. coli* expression host and grown on selective media supplemented with antibiotics, this process takes approximately 24 hours. The next day, DNA of the resulting colonies is extracted and barcodes are ligated using a PCR reaction. The PCR amplified DNA is used as input for the sequencing library process which takes 2 hours. The sequencing library is loaded into the MinION nanopore sequencer and run between 6 and 48 hours. Finally, the obtained sequencing data is demultiplexed, error corrected[21] and annotated using the CARD database[22] in the poreFUME pipeline.

Nanopore sequencing works by threading a DNA molecule through a nanopore embedded in a membrane. When a voltage is applied over the membrane, an ion current is established and this current is modulated when DNA bases pass through the pore. This current-signal depends on the identity of the base that resides in the pore and is, converted into a nucleotide sequence by downstream software. Using the MinION read lengths larger than 100 kb have been reported [17]. The MinION has already been applied to study various aspects related to antibiotic resistance and infection microbiology, including determining the structure and chromosomal insertion site of a bacterial antibiotic resistance island in *Salmonella* Typhi[16], detection of carbapenemases and ESBL genes as well as their position in Gram-negative pathogenic isolates[18] [19], and identification of species and resistance profiles of *Staphylococcus aureus* and *Mycobacterium tuberculosis* isolates[20]. Yet, nanopore sequencing has so far not been applied to profile the resistome of a complex microbial community.

In this study we developed the poreFUME workflow to characterize the resistome of a clinical samples (Figure 1) using nanopore sequencing. Metagenomic expression libraries were constructed using fecal samples from a hospitalized patient as input. The libraries were selected on solid media containing various antibiotics and DNA was extracted from the surviving clones expressing metagenomic inserts conferring antibiotic resistance. The extracted DNA was sequenced using nanopore sequencing. Finally the sequence data was processed using the poreFUME computational pipeline which demultiplexes the barcodes, increases the data quality and annotates antibiotic resistance genes.

## Results

We constructed a metagenomic expression libraries from fecal samples obtained from an Intensive Care Unit (ICU) patient as described in [23] (Materials and methods). The library size ranged between 2.9 – 8.8 x 10 bp of DNA (Supplementary table 1). The metagenomic libraries were plated on solid agar media containing inhibitory concentrations of the antibiotics: tobramycin, spectinomycin, ampicillin, cefotaxime, azithromycin, tetracycline or fosfomycin (Supplementary table 2). Clones from the metagenomic libraries able to tolerate each of these seven different antibiotics were detected in all libraries (Supplementary table 2). From each antibiotic plate a representative number of clones were selected (in total 864), pooled, barcoded using PCR and prepared for sequencing using the MinION nanopore sequencer (Materials and methods).

Nanopore sequencing yielded 95.1 Mbase in 62,890 high-quality ‘passing filter’ two-direction (2D) reads with a mean length of 1,513 bp (library A)(Supplementary Figure 1). As an internal control we multiplexed the sequencing library with 8 other unrelated samples (library B). Due to multiplexing with unrelated samples library B generated only 4,959 sample specific 2D reads (Supplementary table 3). The subsequent part of this study focuses exclusively on the use of high-quality ‘passing filter’ two-direction (2D) reads.

The first step of poreFUME is to demultiplex the barcodes. We identified all the 39 experimentally attached barcodes in both the nanopore sequencing libraries (Figure 2). The abundance showed a significant correlation of log transformed abundance with the Pearson correlation test (R^2^ = 0.75, p< 10^-12^) between the two libraries, highlighting the reproducibility of the sequencing and barcode demultiplexing step. Due to the smaller library size of nanopore library B the remainder of this study focuses on library A.

**Figure 2:**
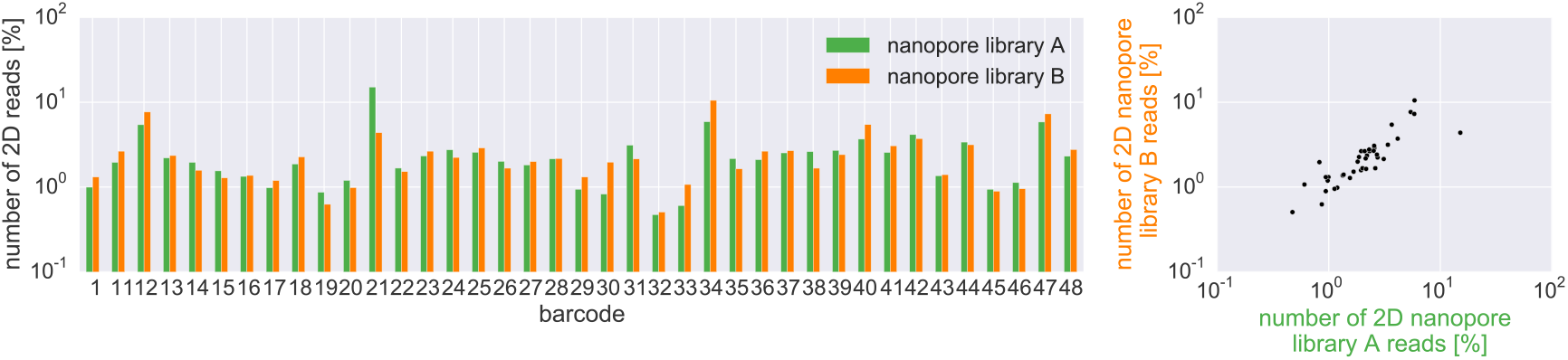
Barcode distribution of the 24,126 2D-reads nanopore library A (green) and control library B (3,361 2D-reads, orange). The Pearson correlation test (R^2^ = 0.75, p< 10^-12^) shows the significance of the log transformed abundance relationship between the two nanopore libraries.

The sequencing data obtained with MinION nanopore R7 chemistry has an 2D read accuracy of ~85% [21]. This relative high error rate can be mitigated using error correction, with tools such as nanocorrect [21]. Nanocorrect is implemented as second step in the poreFUME pipeline. Nanocorrect has been applied previously to increase the nanopore read accuracy from 80.5% to 95.9% [21]. The algorithm identifies overlapping reads using DALIGNER [24] and calculates a consensus sequence, using partial-order alignment (POA) software [25]. Two rounds of error correction where conducted by the poreFUME pipeline.

We annotated the error corrected sequencing data for the presence of antibiotic resistance genes using Comprehensive Antibiotic Research Database (CARD)[22]. Using the CARD database, 26 different antibiotic resistance genes were identified in the nanopore data set (Figure 4). A variety of antibiotic resistance genes were detected with a mean sequence identity of 97.1%, including beta lactamase genes (*CTX, TEM* and *CblA*), genes coding aminoglycoside modifying proteins (from different subclasses of AAC, ANT and APH enzymes) and diverse genes encoding ribosomal and efflux mediated resistance towards tetracycline antibiotic, among others (Figure 3 and S3).

**Figure 3:**
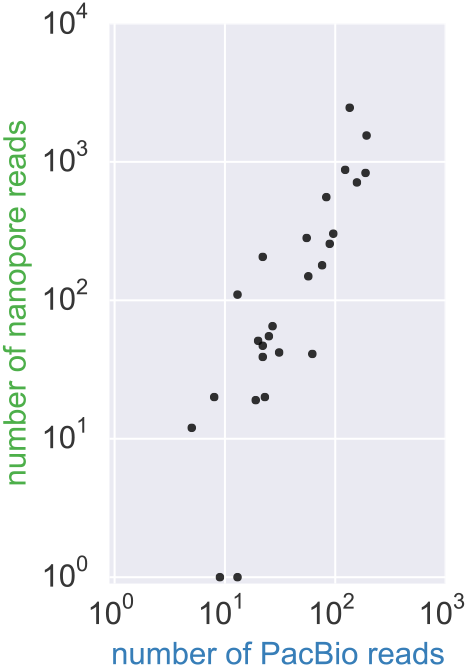
Distribution of the occurrence of the 26 different CARD genes found in the PacBio and nanopore dataset. The Pearson correlation test (R^2^ = 0.71, p< 10^-7^) showed a significant relationship between the nanopore and PacBio dataset as assessed by the log-transformed proportion of CARD hits found. Threshold for CARD identification are gene coverage of >50% and >80% sequence identity.

**Figure 4:**
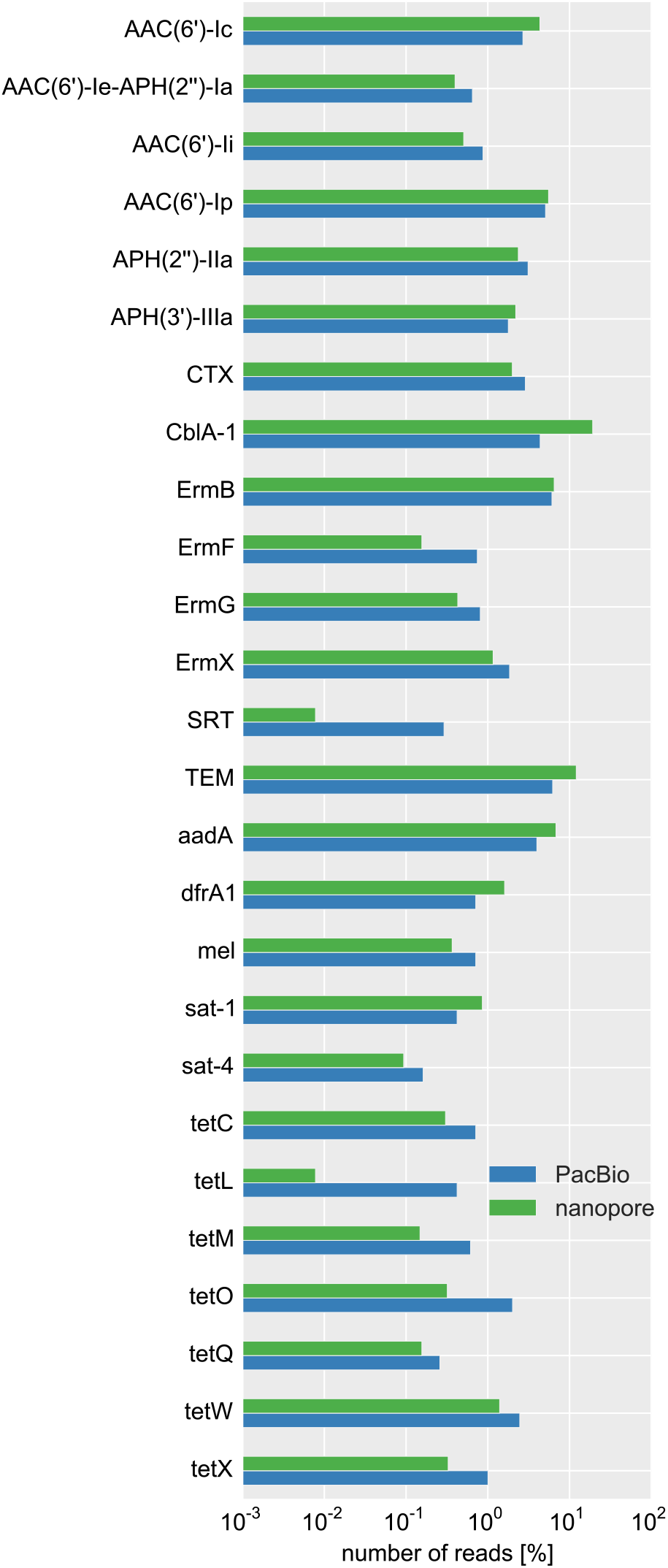
Abundance of 26 CARD genes in the PacBio (blue) and nanopore (green) dataset as show as fraction of the total reads contained in each dataset. The threshold for CARD identification are a gene coverage of >50% and >80% sequence identity.

To validate the nanopore sequencing results, we sequenced the same barcoded DNA using PacBio SMRT technology yielding 93.5 Mbase of DNA in 68,144 reads (with >99% accuracy) from two sequencing cells. After annotation with the CARD database, we observed that the exact same set of 26 antibiotic resistance genes detected in the nanopore dataset were also present in the PacBio dataset (Figure 4). The mean sequence identity of the genes identified in the CARD database is for the PacBio dataset with 97.8% slightly better then that of the nanopore dataset with 97.2% (Supplementary Figure 2). The abundance of reads between the nanopore and PacBio dataset was in good agreement as calculated using the Pearson correlation test (R^2^ = 0.71, p< 10^-7^) (Figure 3).

To further test the accuracy of our nanopore data set we sequenced the selected libraries using Sanger sequencing. The sequence identity between the Sanger reads and the non-corrected 2D reads of nanopore library A was 85.8% (Supplementary Figure 2), this confirms the higher error rate of the used MinION reads [16, 21]. However by using two rounds of error correction implemented by poreFUME the sequence identity of the nanopore reads was improved from 86.8% to 97.8%, which enables accurate resistome mapping using CARD.

**Figure 5:**
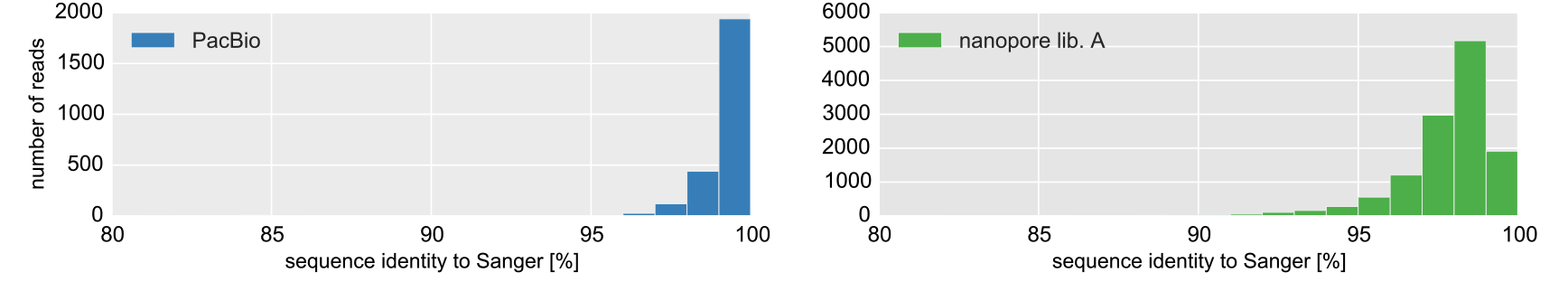
Histogram of the sequence identity of the 2D nanopore sequence reads of library A (12,820 reads) after two rounds of error correction, showing a mean sequence identity to the Sanger data set of 97.8%. The PacBio dataset after a single round of consensus calling using pbdagcon (3,086 reads) shows a mean sequence identity to the Sanger dataset of 99.3%

Nanopore reads are longer and typically capture the entire metagenomic insert. In this way analysis of the context of the antibiotic resistance gene is simplified. As an example to link genotype and phenotype we investigated the 244 nanopore reads that were selected on plates containing spectinomycin and resulted in a CARD annotation. All the 244 reads contain the *aadA* gene which encodes an aminoglycoside nucleotidyltransferase known to confer resistance to spectinomycin (Figure 6a). In 74 reads *aadA* was flanked by *sat-1* which encodes a streptothricin acetyltransferase and confers resistance to streptothricin, again *sat-1* is likely to be co-selected with *aadA*. In 44 reads the *aadA* was the only gene detected, however in 126 reads *aadA* was flanked by *dfrA1.* The *dfrA1* gene confers resistance to trimethoprim and is not known to confer resistance to spectinomycin. Alignment of the longest nanopore read containing both *dfrA1* and *aadA* against the NT database showed that the two highest scoring hits are part of an integron class I (Figure 6b). Nanopore sequencing is thus able to identify such differing contexts of antibiotic resistance genes, which can impact the probability of a pathogen to acquire specific antibiotic resistance genes[26].

**Figure 6a:**
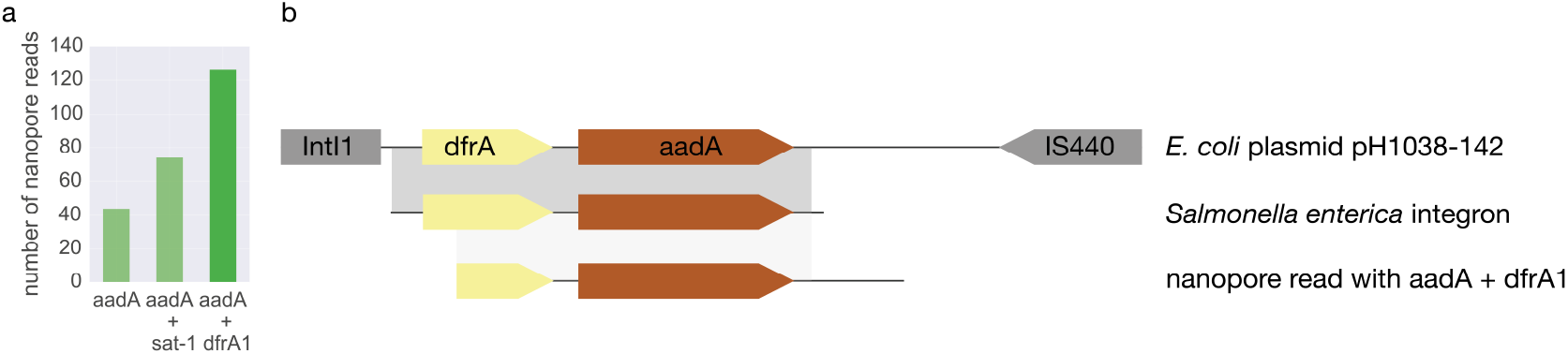
Example of CARD genes found in the nanopore dataset that were plated on spectinomycin. Of the 244 nanopore reads that were recovered on spectinomycin, all contain the *aadA* which is an aminoglycoside nucleotidyltransferase known to confer resistance to spectinomycin. In 126 reads *aadA* is flanked by *dfrA1* which confers resistance to trimethoprim and in 74 reads *aadA* is flanked by *sat-1* which confers resistance to streptothricin. The genes *sat-1* and *dfrA1* that do not confer known resistance to spectinomycin are thus co-selected with *aadA*. b: Alignment against the NT database of the longest corrected nanopore read of the 126 reads containing *aadA* and *dfrA* obtained from functional selection on spectinomycin. The corrected nanopore read shares 99% sequence identify with the two top scoring hits in the NT database: (light gray) the class I integron from *Salmonella enterica* subsp. enterica serovar (genbank: HQ132378.1) and the *E. coli* plasmid pH1038-142 (genbank: KJ484634) described by Wang et al.[27]. *IntI* denotes integron class I, *IS440* an transposon, *dfrA* encodes a dihydrofolate reductase known to confer resistance against trimethoprim and aadA is an aminoglycoside nucleotidyltransferase known to confer resistance to spectinomycin

## Discussion & Conclusion

Hospital settings, including ICUs, are hotspots for the emergence and selection for antibiotic resistant organisms. In this study we successfully applied nanopore sequencing to characterize the gut resistome of an ICU patient using metagenomic functional selections. Nanopore sequencing is known to have a higher error rate compared to other sequencing technologies; however, implementing a double error correction scheme in poreFUME we achieve accuracies above 97%, which enables reliable resistance gene annotation and comparable results to that of PacBio SMRT sequencing. In this study both sequencing platforms enabled the reliable identification of 26 unique antibiotic resistance genes. This, along with the rapid turnaround time of the poreFUME workflow, suggests that it could be applied as a possible resistome monitoring tool.

Only 39 % of the nanopore 2D reads had their barcodes successfully demultiplexed in this study. Improvements to the protocol can be made by making use of barcodes with a larger editing distance. In accordance, the currently used PacBio barcodes are deprecated and replaced by a set with a larger editing distance, which allows for better separation and a higher barcode identification rate. However, even the 61 % loss due to barcode demultiplexing does not hamper downstream resistome analysis using poreFUME.

In conclusion, the poreFUME pipeline provides a promising alternative to other next-generation sequencing alternatives [6, 15] and can be used to rapidly profile the resistome of both environmental and gut microbial communities [4-6]. We foresee that rapid resistome profiling tools such as poreFUME could aid the implementation of personalized antibiotic treatment in high risk patients.

## Material and Methods

### Experimental work

#### Ethics statement

Ethical approval for this study was obtained from the institutional review board of the University Medical Center Utrecht (Utrecht, The Netherlands). During hospitalization informed consent was waived. The collection of fecal samples after hospitalization was done with the subject written consent.

#### Sample collection

Longitudinal fecal samples were collected from a human adult who received ICU care at the Utrecht University Hospital in Netherland. The subject was a patient who after surgical intervention was admitted to ICU. Five longitudinal samples were collected upon admission, during ICU stay and 8 months after hospitalization. During the recovery at ICU, the patient received antibiotics from at least the cephalosporin group (beta-lactam) and SDD (Selective Digestive Decontamination), which included the aminoglycoside (tobramycin) and polymyxin (colistin) antibiotic class.

#### DNA extraction

DNA was obtained from Utrecht University Hospital in the Netherlands. The DNA extraction method was described previously [28].

#### Construction of metagenomic libraries for functional selections

Construction of metagenomic libraries was done following the protocol [23] with slight modifications. DNA was sheared using a Covaris shearer to an average size of 2 kb. DNA was blunt-ended and phosphorylated using an End-Repair kit (Epicentre, USA). Sheared DNA (138 μl) was mixed with 20 μl 10X End Repair buffer, 20 μl 2.5 mM dNTP, 20 μl 10 mM ATP and 2 μl of End-It enzyme. After incubation at 22 °C for 55 min, the end-repair reaction was heat inactivated at 70 °C for 20 min. End-repaired DNA was size selected by electrophoresis. Agarose gel slices selected from the size range 0.5 – 5 kb were purified using the Gel Purification Kit (Fermentas). Gel purified DNA was ligated into vector pZE21-MCS-1 [29] using Fast-link DNA ligase (Epicentre). For this purpose, the concentration of gel-purified and end-repaired DNA insert was adjusted to 200 ng/μl. Ligation reaction was set up with 2.5 μl of DNA inserts (200 ng/μl), 0.5 μl 10X Ligation buffer, 0.25 μl 10mM ATP, 0.25 μl dH_2_O, 0.5 μl HincII cut pZE21-MCS-1 vector (100 ng/μl). The ligation mixture was incubated at 22 °C for 16 h and finally heat inactivated at 70 °C for 20 min. Three μl of ligation reaction was electroporated into 50 μl electrocompetent *Escherichia coli* Top10 cells (Invitrogen). After electroporation in a 2 mm cuvette (2000 V, 25 μF, 200R), cells were recovered in 1 ml SOC medium for 1 h at 37 °C. Determination of the transformation efficiency was done by plating out 1 μl and 0.01 μl of recovered cells onto LB agar plates containing 50 μg/ml kanamycin (pZE21-MCS-1 vector contains a selectable marker for kanamycin resistance) [29]. Colony forming units (CFU) were counted after overnight incubation at 37 °C. For each library, the insert size distribution was estimated by gel electrophoresis of the PCR products obtained by amplifying the insert using primers flanking the HincII site of the multiple cloning site of the pZE21-MCS-1 vector (>pZE21_81_104_57C and pZE21_151_174rc_58C; Supplementary table 4) [23].

The size of each of the metagenomic libraries for functional selection was determined by multiplying the average PCR based insert size with the number of colony forming units (CFU). The size of the 5 metagenomic libraries for functional selection is listed in Supplementary table 1. The rest of the recovered cells after transformation was inoculated into 10 ml of LB broth supplemented with 50 μg/ml kanamycin and grown overnight at 37 °C, 180 rpm. The overnight cultures were stored with 15 % glycerol at -80 °C.

#### Functional selection of antibiotic resistance clones

The overnight cultures grown and stored at -80 °C allowed each clone after transformation to amplify (e.g. total cell count would increase from 5x10^5^ CFU containing the plasmid per ml to 5x10^8^ CFU/ml after overnight incubation and storage at -80°C). Resulting amplification of the particular clone in the library was taken resolved by plating each library approximately 100X coverage. That is, each unique clone in the library was screened by plating out approximately 100 copies. For each library, clones carrying antibiotic resistance determinants were selected by plating onto solid LB agar supplemented with one of the seven antibiotics: tobramycin, spectinomycin, ampicillin, cefotaxime, azithromycin, tetracycline or fosfomycin at concentrations that were inhibiting the wild type strain (Supplementary table 2). The CFU was determined after overnight incubation at 37 °C (Supplementary table 2).

#### Sequencing of antibiotic resistance clones

From each antibiotic plate, a representative number of clones were selected for sequencing and further analysis of antibiotic resistant genes. Singe colonies selected on antibiotic resistance plates were picked up into 96-well plates (each 96 well contained 200 μl LB broth supplemented with 50 μg/ml kanamycin). The selected clones were grown overnight at 37 °C. The clones were transferred using a 96-pin replicator into new 96-well plates and onto squared LB agar plates supplemented with 50 μg/ml kanamycin. Singe clones from 96-well plates were Sanger sequenced using primers listed in table Supplementary table 4 by Beckman Genomics, UK.

The clones from the individual square agar plates were collected by adding 5 ml dH_2_0 and scraped off with an L-shaped cell scraper. The washing step was repeated twice to remove all the cells from the plate. The bacterial cells were then pelleted by centrifugation at 5.000 rpm x g for 10 min. The supernatant was discarded and the pellet was dissolved in 10 ml of dH_2_O. Two ml of the collected bacterial cells was used for plasmid extractions with the Plasmid Mini Kit (Invitrogen). The rest of the cells were heat inactivated at 95 °C for 10 min and stored as raw bacterial cell pellet. For nanopore and PacBio sequencing, primers were synthetized that amplify the common region on pZE21-MCS-together with the specific barcodes from PacBio (Supplementary table 5). One ng of DNA or 1 μl of raw bacterial cell pellet was amplified by PCR. Amplified and barcoded DNA was size selected by electrophoresis. Agarose gel slices selected from the size range 1 – 5 kb were purified using Gel Purification Kit (Fermentas). In total 39 barcodes (1 and 11-48) were multiplexed.

#### Nanopore sequencing library preparation

The nanopore sequencing library B was prepared during poreCamp as briefly described below. DNA QC was performed using Qubit dsDNA High Sensitivity Assay Kit (Q32851, Thermo Fisher Scientific, USA) and 2200 TapeStation (G2964AA, Agilent, USA). Sequencing library preparation was carried out with Nanopore Genomic Sequencing Kit SQK-MAP006 (Oxford Nanopore, UK) and a PCR free ‘native barcoding’ kit according to the manufacturer’s protocol. The NEBNext Ultra II End Repair/dA Tailing module (E7546S, NEB, USA) was used to prepare 1000 ng of the functionally selected DNA. End-prepared DNA was ligated with native barcode adapters NB04 from Oxford Nanopore using Blunt/TA Ligase Master Mix (M0367S, NEB, USA). The resulting DNA was pooled with 8 other unrelated barcoded libraries by equivalent weight. The pooled sample was mixed with the ‘Native Barcoding Adapter Mix (BAM)’ and ‘Native Barcoding Hairpin Adapter (BHP)’ together with Blunt/TA Ligase Master Mix (M0367S, NEB, USA), and after incubation HP tether was added. The reaction mixture was cleaned up using prewashed Dynabeads MyOne Streptavidin C1 (65001; Thermo Fisher Scientific, USA).

The sequencing library A was prepared using the same protocol as library B, but the barcoding and pooling steps were omitted.

#### Nanopore sequencing

The MinION was primed twice for 10 minutes with 500 μl priming solution (250 μl nuclease free water, 237 μl 2x Running Buffer, 13 μl Fuel Mix). For sequencing, 6 μl library was mixed with 65 μl nuclease free water, 75 μl 2x Running Buffer and 4 μl Fuel Mix (SQK-MAP006, Oxford Nanopore, UK) and immediately loaded to a MinION. The ‘SQK-MAP006 Scripts for Yield Monitoring Switch, Bias-Voltage Remux Tuning & Pore Shepherding’ by John Tyson (personal communication) were used in the MinKNOW software to sequence the library.

### Data analysis

#### Nanopore data processing

The sequencing data was basecalled using Metrichor. The Metrichor workflow for sequencing library A included additional native barcode demultiplexing. Poretools[30] was used to extract 2D reads using the *poretools fasta --type 2D* command. Next the 2D reads were analyzed using poreFUME.

#### poreFUME nanopore sequence analysis

The poreFUME pipeline consists of three steps. First, the reads are demultiplexed on barcode using the Smith-Waterman algorithm[31]. Barcodes are detected within 60 (Library A) or 120 (Library B) basepairs of the read ends. Barcode alignment was scored using +2.7 for match, -4.5 for mismatch, -4.7 gap opening and -1.6 for gap extension. A score threshold of >58 was used for the combined score of the asymmetric barcodes. Second, the demultiplexed reads were error corrected using the original nanocorrect protocol [21]. The original nanocorrect protocol implements a minimum read coverage of 3x, to ensure that only high-quality data will be outputted. Since we were also interested maximizing sequence diversity, we adjusted the minimum coverage to 1x in the second round of nanocorrect by modifying the *min_coverage* parameter from 3 to 1. In the last step of poreFUME the error corrected reads were mapped against the CARD database[22] using blastn[32] (version 2.4.0) with the parameters *max_hsps* 1 and *max_target_seqs* 1000. Closely related genes such as *TEM, CTX, MIR* and *SRT* (ie. *SRT-1* and *SRT-2*) were masked and only reported as such (ie. *SRT*). For each individual read the BLAST hits were sorted by bitscore and the highest scoring CARD hit on each segment was kept. For successful CARD gene calling two threshold were set: a sequence identity of >80% and a >50% coverage of the original gene in the CARD database.

#### Sanger sequence data analysis

The Sanger sequencing resulted in 770 ‘forward’ and 779 ‘reverse’ sequencing reads. Sanger DNA sequences were imported to CLC Genomic Workbench (version 7.6.4). Sequences were vector and quality trimmed (Q 0.01) and assembled using the ‘Assemble Sequences’ module. Contigs with a length of <500 basepairs were omitted from further analysis.

#### PacBio data analysis

PacBio sequences were obtained from the Norwegian Sequencing Centre at the University of Oslo in two flowcells on the Pacific Biosciences RS II instrument using P6-C4 chemistry. The metrics for the total set are listed in Supplementary table 6. Raw PacBio data from the flowcells was analyzed with PacBio SMRT^®^ Portal version 2.3.0 and reads were extracted using the RS_ReadsOfInsert protocol (version 2.3.0). The RS_ReadsOfInsert protocol was run with a minimum predicted accuracy of 99, and minimum read length of insert length of 100 bp. Additionally paired-end barcode detection was performed using the pacbio_barcodes_paired scheme containing 48 unique barcode pairs. A minimum barcode score of 15 was used in both cells. The extracted reads of insert were grouped by individual barcode and exported in the fastq. The final yield is reported in Supplementary table 5. PacBio reads were collapsed with Pbdagcon (https://github.com/PacificBiosciences/pbdagcon, version f19aed1) using dazcon with the flags --*only-proper-overlaps* and --*coverage-sort* and parameter --*min-coverage* 0.

## Declarations

### Availability of data and materials

The source code for poreFUME is available at https://github.com/EvdH0/poreFUME and the full analysis of the data is available at https://github.com/EvdH0/poreFUME_paper/. The PacBio and nanopore sequence data is deposited at ENA with project code PRJEB14994.

## Acknowledgement

This research was funded by the EU FP7-Health Program Evotar (282004), The Novo Nordisk Foundation and The Lundbeck Foundation. Additionally this work was supported by the European Union Seventh Framework Programma – INT FP/7/2012/317058 to EvdH. MMHE acknowledge the help and assistance of Ali Syed and other members of computerome, Danish National Supercomputer for Life Sciences, in using the HPC facilities. Library B was prepared during the porecamp MinION workshop in Birmingham December 2015, all participants and organizers (Nick Loman, Matt Loose, Mick Watson, Josh Quick, John Tyson and Justin O’Grady) are thanked, especially Kathrin Lang for help with writing up of the sequencing protocol. The PacBio sequencing service was provided by the Norwegian Sequencing Centre (www.sequencing.uio.no), a national technology platform hosted by the University of Oslo and supported by the “Functional Genomics” and “Infrastructure” programs of the Research Council of Norway and the Southeastern Regional Health Authorities.

## Author contributions

EvdH and MOAS conceived the study. WvS obtained IRB approval and organized fecal sample collection. LI designed the experiments, carried out the metagenomic library construction, functional selections and obtained Sanger sequencing data. MMHE provided input on the poreFUME pipeline and provide assistance with an early prototype. EvdH carried out the nanopore sequencing library preparation together with AK. EvdH ran the sequencing experiments and wrote the poreFUME code. EvdH wrote the manuscript with input from the other authors.

## Supplementary information

**Supplementary table 1:** Library size in basepair of the constructed metagenomic libraries.

| Sample | Size (bp) |
| --- | --- |
| 120A | 288120000 |
| 120B | 546040000 |
| 120C | 881280000 |
| 120D | 574560000 |
| 120E | 479360000 |

**Supplementary table 2:** inhibitory concentrations of antibiotics used and the number of clones per agar.

| Antibiotic | Abbreviation | Concentration<br>(µg/ml) | Sample (n clones / agar plate) |  |  |  |  |  |
| --- | --- | --- | --- | --- | --- | --- | --- | --- |
|  |  |  | 120A | 120B | 120C | 120D | 120E | total |
| Tobramycin | TOB | 6 | 121 | 216 | 480 | 950 | 240 | 2007 |
| Spectinomycin | SPC | 32 | 220 | 180 | 640 | 360 | 184 | 1584 |
| Ampicillin | AMP | 64 | 156 | 180 | 140 | 524 | 216 | 1216 |
| Cefotaxime | CFT | 2 | 48 | 58 | 14 | 6 | 139 | 256 |
| Azithromycin | AZY | 16 | 26 | 156 | 154 | 640 | 36 | 1012 |
| Tetracycline | TET | 6 | 6 | 142 | 100 | 71 | 15 | 334 |
| Fosfomycin | FOS | 128 | 58 | 124 | 288 | 111 | 52 | 633 |

**Supplementary table 3:** Statistics of the Sanger, PacBio and two nanopore sequencing sets.

| sample | workflow | number<br>of<br>entities | mean<br>length<br>[bp] | yield<br>[Mbase] | mean<br>sequence<br>identity of<br>entities to<br>raw<br>Sanger<br>reads [%] | mean<br>sequence<br>identity of<br>hits to<br>CARD<br>database<br>[%] |
| --- | --- | --- | --- | --- | --- | --- |
| Sanger | raw | 1,568 | 952 | 1.5 |  | 98.76 |
|  | assembled | 111 | 1,532 | 0.2 | 99.96 | 98.87 |
| PacBio | raw | 68,144 | 1,370 | 93.4 | 99.77 | 98.64 |
|  | assembled | 3,086 | 1,626 | 5.0 | 99.28 | 97.83 |
| nanopore<br>library A | raw 2D | 62,890 | 1,513 | 95.1 | 85.93 | 85.80 |
|  | after debarcoding | 24,126 | 1,440 | 34.7 | 86.81 | 86.41 |
|  | after nanocorrect round 1 | 12,820 | 1,541 | 19.8 | 96.88 | 96.19 |
|  | after nanocorrect round 2 | 12,820 | 1,540 | 19.7 | 97.76 | 97.08 |
| nanopore<br>library B | raw 2D | 4,959 | 1,587 | 7.9 | 86.15 | 85.65 |
|  | after debarcoding | 3,361 | 1,426 | 4.8 | 86.38 | 85.80 |
|  | after nanocorrect round 1 | 2,392 | 1,444 | 3.5 | 96.81 | 96.34 |
|  | after nanocorrect round 2 | 2,392 | 1,441 | 3.4 | 97.59 | 97.19 |

**Supplementary table 4:**
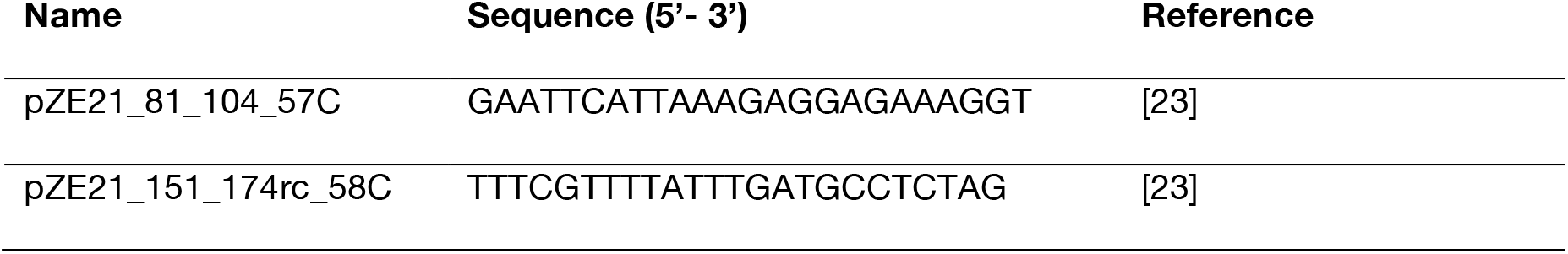
Primers used in this study for Sanger sequencing.

**Supplementary table 5:**Primers used to barcode the sample for PacBio and nanopore sequencing

Attached csv file

**Supplementary table 6:**
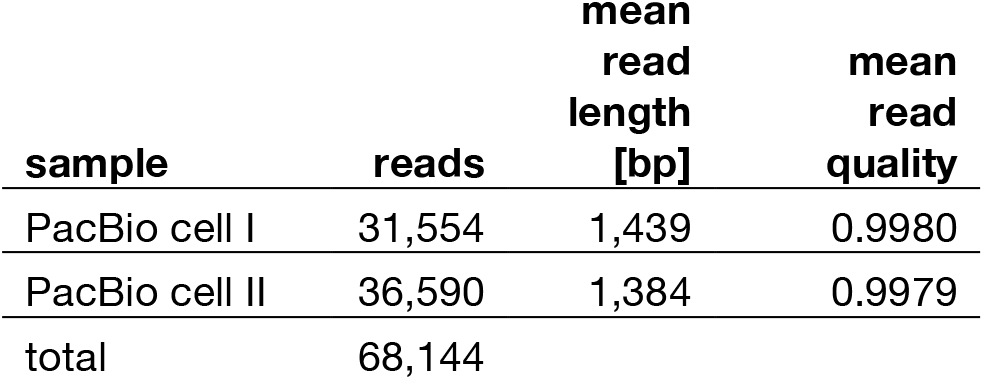
Statistics of the two sequencing cells of the filtered (quality >99% and barcode demultiplexed) PacBio reads as reported by SMRT Portal. Both PacBio cells were used in the analysis.

| **sample** | **reads** | **mean**<br>**read**<br>**length**<br>**[bp]** | **mean**<br>**read**<br>**quality** |
| --- | --- | --- | --- |
| PacBio cell I | 31,554 | 1,439 | 0.9980 |
| PacBio cell II | 36,590 | 1,384 | 0.9979 |
| total | 68,144 |  |  |

**Supplementary Figure 1:**
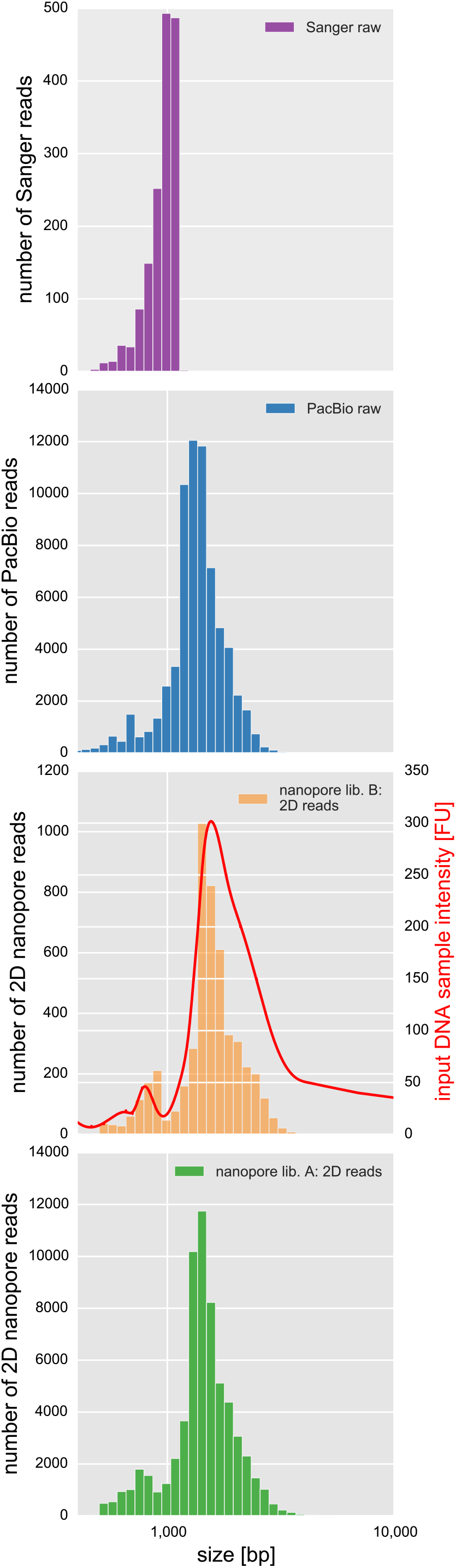
Read length distribution of the control Sanger (purple) data with a mean of 952 bp and PacBio (blue) data with a mean read length of 1370 bp. The 2D read length of nanopore library A (orange) with a mean of 1587 bp is overlaid with the DNA input sample intensity (red) measured using a TapeStation showing agreement between the length distribution. Nanopore library B has a mean read length of 1513 bp.

**Supplementary Figure 2:**
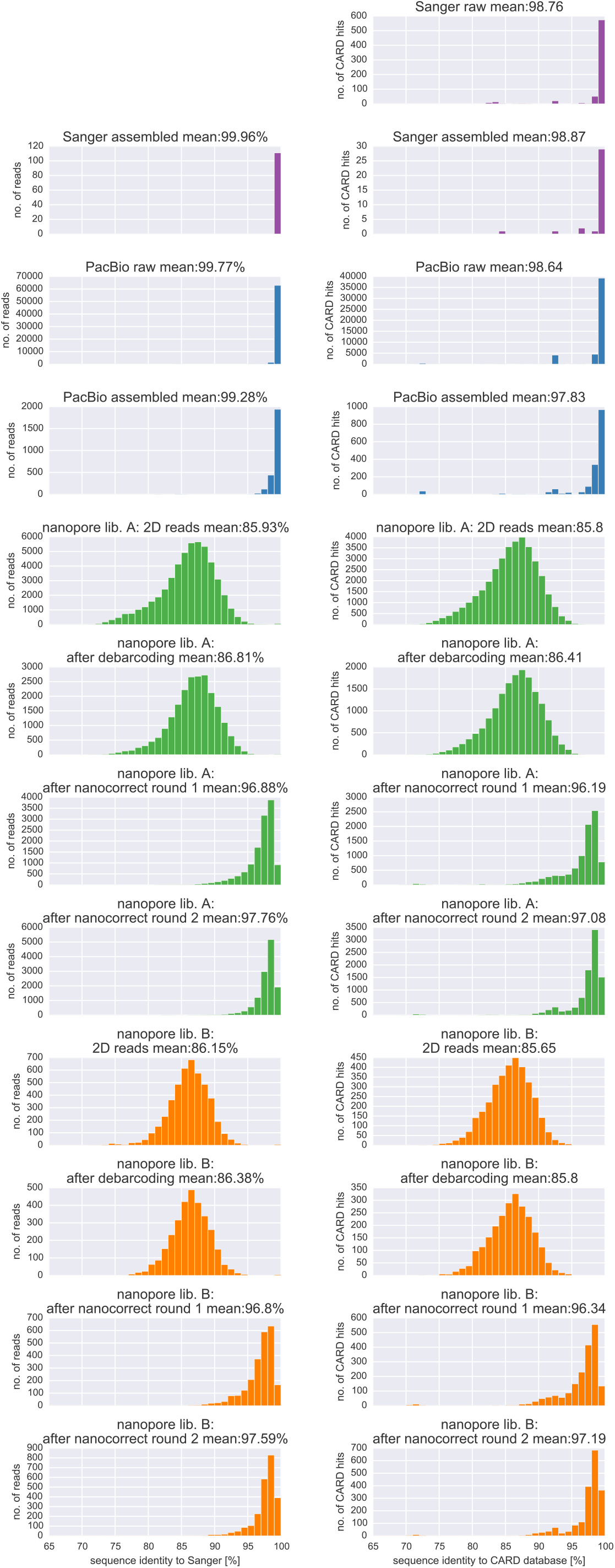
Sequence identity of the Sanger (purple), PacBio (blue), nanopore library A (orange) and nanopore library B (green) dataset measured against the Sanger dataset in the left column (with >500 bp in alignment length). The sequence identity of the highest scoring hits (with a gene coverage requirement of >50%) in the CARD database are shown in the right column.

**Supplementary Figure 3:**
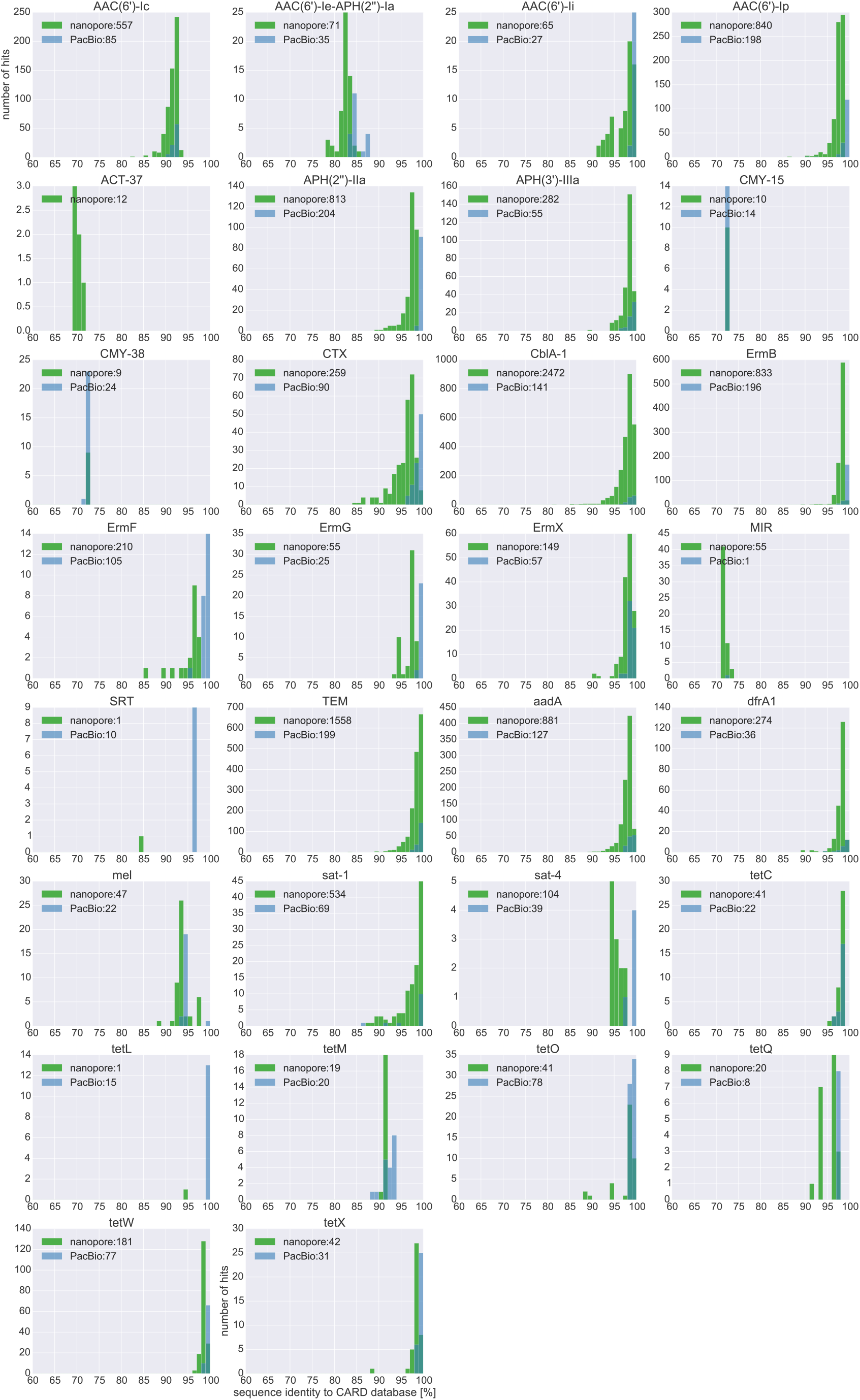
Overview of unique antibiotic resistance genes of the CARD database that are found in the nanopore library A (green) and PacBio set (blue). The number of reads over the sequence identity [%] to the respective gene in the CARD database is shown. In the legend box the total number of reads for each dataset is shown.

